## Supplemental Information for "Communication with Surprise – Computational Principles of Goal Signaling in Novel Human Interactions"

#### Supplement Text and Figures:

##### Original and Modified Tacit Communication Game

The original version of the TCG[1–4] featured two objectives: the Receiver's goal location and target orientation, displayed in Figure S1a. However, the former resulted in an almost unanimous use of the "pause" strategy (pausing at the Receiver's goal state), lacking any experimental variability in the message trajectories, whereas the second goal elicited very diverse strategies. We decided to exclusively focus on the location problem and aimed to generate more diverse spatial trajectories. We therefore made the following changes to the original task:

1. *Targets with the same geometrical objects*. To exclude the orientation problem, we used the same objects in different colors for both players as their goal configuration as displayed on the modified grid on Figure S1b, and instructed the participants that finding the correct goal locations was the only objective of the game.
2. *Increase the grid size.* We modified a 3×3 grid to a 4×4 grid to introduce more complexity in the game by including more goal configuration.
3. *No real-time display of the Sender's message*. When the Sender first moves her game token, the Receiver does not see the message. Afterward, the message is played back to both participants with a 1.2 s duration for each step. This does not preclude the Sender from using the "pause" strategy, but it makes it obsolete because the Receiver will not see it during the replay. Thus, this change encourages the Sender to create a more comprehensive range of behavioral patterns.

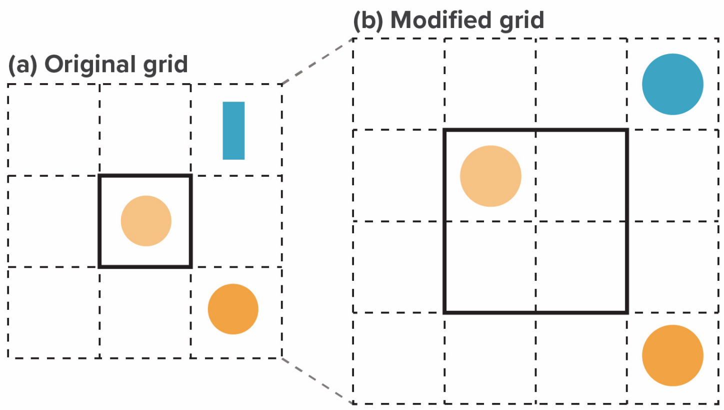


**Supplementary Figure S1 | Original and Modified Version of the TCG**
(a) The Original 3x3 grid with different shapes of tokens for Sender and the Receiver. (b) The modified game setup involved using identical geometric objects in different colors for both players as target goals, displayed on a modified 4×4 grid, with the sole objective of finding the correct goal locations. This change, along with the enlargement of the grid from 3×3 to 4×4, was implemented to increase the game's complexity through a greater variety of goal configurations.

#### Belief-Based Model

The BBM stems from the foundational principles of the Simulation Theory of Mind, enabling computational agents to predict their partner's choices by presuming they think similarly[4]. Its core computational unit is a belief distribution over all states that encodes the most likely location of the Receiver's goal state. The model calculates the set of all possible goal locations and within each goal configuration all possible messages (L, M) and selects the one with the highest belief probability at the Receiver's goal state.

During a trial:

1. The Sender observes the Sender's and Receiver's goal location (ls, lr) and selects a message (m) from all potential messages (M) to send to the Receiver.
2. Upon receiving the message, the Receiver identifies the location (l), as his goal location.
3. Success is achieved if the chosen locations align with the true goal configuration.

Initially, the sender establishes a uniform belief distribution across the states of all the messages (M). As interactions unfold, this distribution is continually updated and recalibrated, influenced by the Receiver's behavior and the Sender's inferred depth of the Theory of Mind. Depending on the agent's depth of Theory of Mind, the resulting behavior varies:

- Zero-order theory of mind (ToM-0): Agents with ToM-0 cannot reason about the mental content of others. They try to find actions that randomly would lead to both agents matching their tokens to the goal configuration. Thus, the Receiver would randomly select his goal state among the states of the Sender's message.
- First-order theory of mind (ToM-1): Agents with ToM-1 not only consider their own goals, but also try to understand their partner's goals. When the Sender operates with a first-order Theory of Mind, they recognize that sending the same message could lead

the Receiver to take an alternative action if the previous trial was not successful. Consequently, while adjusting the belief distributions for that message, the Sender eliminates past locations and enhances the belief probabilities of message states that were not selected previously. If the message was successful, then the sender sets the probability of 1 to the chosen location (Receiver's goal location) and zero to the unselected location.

- Higher-order Theory of Mind (ToM-k): Agents with ToM-k, beyond first order, consider the possibility that their partner is also reasoning about their perspective. This recursive reasoning allows them to understand that their partner might be thinking about their thoughts and so on. For instance, a Receiver with a ToM-2 perspective possesses a ToM-1 understanding of the Sender, granting them insight into her reasoning process as previously described.

For a detailed mathematical description of the model, please refer to de Weerd at al[4].
Let's look at an example trial illustrated in Figure S2. First, the Sender observes both her own and the Receiver's goal location and then selects a message from the set of all possible messages (a subset of these is represented by a solid line in Figure S2a) to convey to the Receiver. Once the message is received, the Receiver selects a location, highlighted by the red tile in Figure S2c, as the goal location.

Initially, the Sender establishes a uniform belief distribution across the states of the message, as depicted in Figure S2b. This distribution is then modified, in the following trials, based on the Receiver's actions and the presumed depth of Theory of Mind that the Sender operates on.

Figure S2d illustrates a ToM-1 Sender's process of updating belief distributions after observing the Receiver's action (in this case, action was incorrect). If operating under a first-order ToM, the Sender discerns that sending an identical message would lead to the Receiver opting for a different action, given the prior trial's action resulted in failure. Hence, when updating belief distributions for this message, the Sender ascribes a zero probability to the previous action and increases the probabilities of the actions not previously chosen, as shown in Figure S2d.

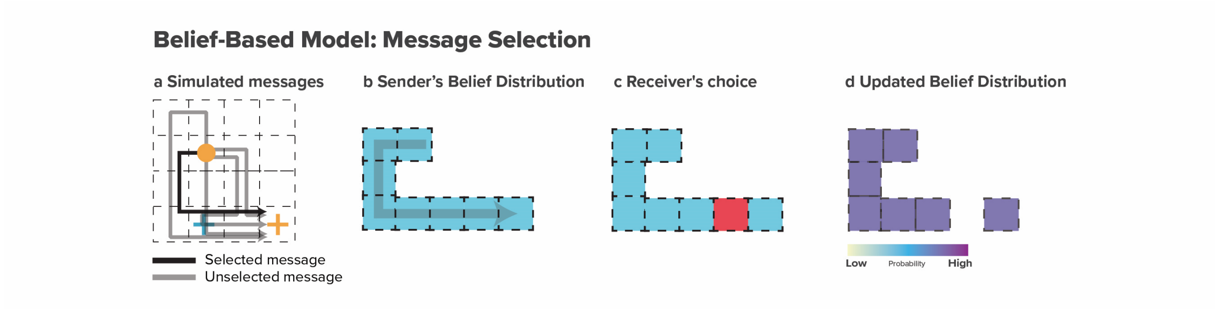


**Supplementary Figure S2 | Model of Belief-Based Communication**
**(a) BBM Simulated Messages:** The Grid showcases varied BBM-generated messages, with one message emphasized by a solid black line. **(b) Sender's Belief:** After message selection, belief probabilities are uniformly distributed across all states. **(c) Receiver's Choice:** Receiver's goal state is randomly selected, highlighted in red. **(d) Updated Belief Distributions:** Sender recalibrates belief distributions considering the receiver's actions and ToM level.

In assessing various models for our study, we carefully considered the operational mechanisms and compatibility of each with empirical data. Our decision to exclude the Belief-Based Model (BBM) from the main model comparisons is grounded in several key considerations.

First, the BBM diverges from other models like the Surprise models in its core mechanism. While the Surprise model emphasizes message creation, the BBM focuses on message selection. This involves defining a belief distribution over possible states on the game board and selecting messages through an exhaustive search for each goal configuration. This process, crucially, demands extensive memory for storing and retrieving belief representations, a requirement that is at odds with the brain's preference for cognitive efficiency. Such an exhaustive and resource-intensive approach seems unlikely as a brain mechanism, given this preference. Second, the BBM operates on complete messages, which impairs its ability to support a step-by-step analysis of message components. This limitation poses significant challenges in aligning the BBM with behavioral and neural data, which often necessitate a more nuanced and detailed examination. Lastly, a critical drawback of the BBM is its lack of free parameters. The absence of adjustable parameters restricts the model's flexibility, making it difficult to fit and compare it effectively with other models.

**Posterior Predictive Check with BBM**

Here we compare message creation by the Surprise model and the BBM with the messages generated by the participants. We employed a Bayesian approach[5] to assess the support for two hypotheses: the null hypothesis ($H_0$), which suggests no difference between the messages generated by each model and the empirical data, and the alternative hypothesis ($H_1$), which posits substantial differences between the model and the empirical data. We quantified the strength of this evidence for or against these hypotheses using a Bayes Factor ($BF_{01}$) for each behavioral index In Figures S3 and S4, asterisks highlight where the model-generated indices diverge from the behavioral data (indicated by a Bayes Factor, BF01, less than 1), and the Surprise Model's illustrations correspond to those presented in the primary text.

**Evidence against BBM.** In contrast to the Surprise model, when comparing the means of human-created message types to the means of the BBM, all the calculated $BF_{01}$ favor the alternative hypothesis, indicating that messages generated by the BBM are different from the messages generated by our participants, suggesting that this model is not capturing human message generation accurately.

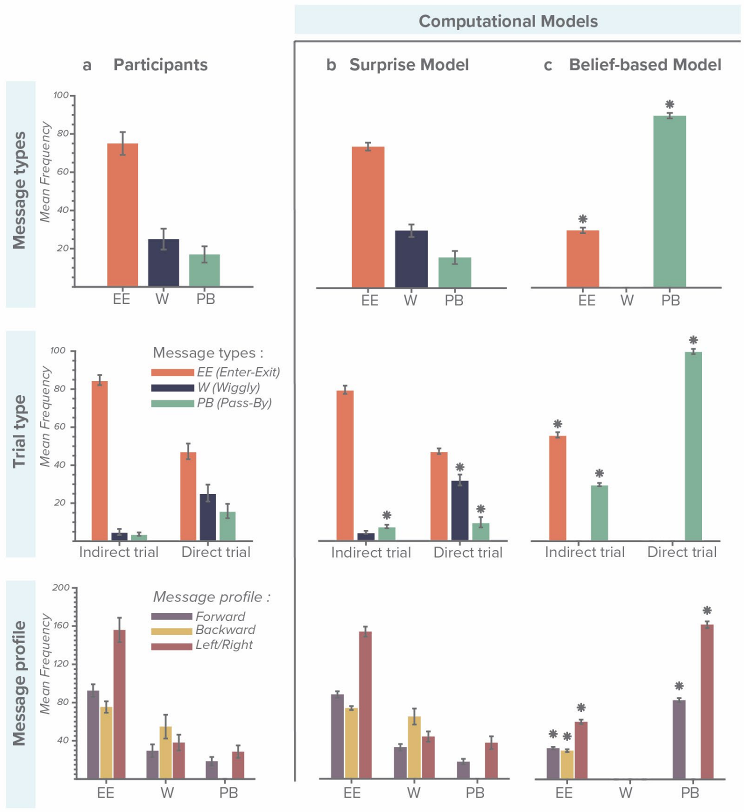


**Supplementary Figure S3 | Comparing Participant-Generated and Model-Simulated Messages for the Dataset 1 (a) Participants:** Participants behavioral data categorized by Message types, Trial types, and Message profile**. (b) Surprise Model:** Surprise model simulated data after model fitting exhibits similarity to participants data ($BF_{01}>1$) **(c) Belief-Based Model:** BBM simulated data reveals differences from participant's data ($BF_{01}<1$*). All error bars represent means +/- SEM.

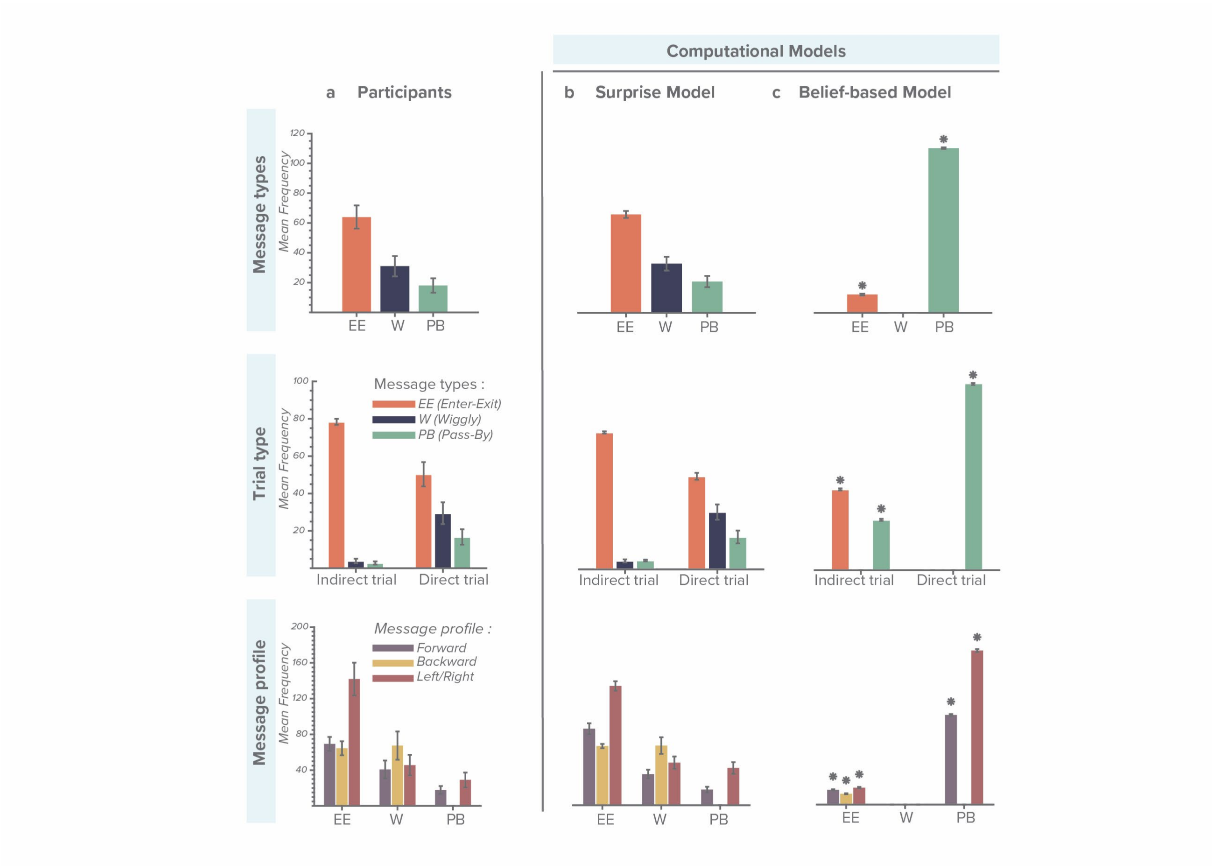


**Supplementary Figure S4 | Comparing Participant-Generated and Model-Simulated Messages for the Dataset 2 a) Participants:** Participants' behavioral data categorized by Message types, Trial types, and Message profile. **(b) Surprise Model:** Surprise model simulated data after model fitting exhibits similarity to participants data *($BF_{01}>1$)* **(c) Belief-Based Model:** BBM simulated data reveals differences from participant's data *($BF_{01}<1$*)*. All error bars represent means +/- SEM.

#### Model Fitting and Model Comparison for Belief-Based Model

Due to the nature of the BBM, which lacks estimable parameters and precludes the calculation of the model likelihood as a cost function, we did not include the BBM in a formal model's selection.

### Supplement Tables:

***Data set 1***

***a***

| | Participants | | SM Model | | State Model | | Movement Model | | $BF_{01}$ | | |
|---|---|---|---|---|---|---|---|---|---|---|---|
| *Message type* | *M* | *SE* | *M* | *SE* | *M* | *SE* | *M* | *SE* | *Participants vs SM Model* | *Participants vs S Model* | *Participants vs M Model* |
| Enter-exit | 63.82 | 5.00 | 61.70 | 1.72 | 26.46 | 7.46 | 12.07 | 0.54 | 10.20 | 6.37E-07 | 1.46E-13 |
| Wiggly | 21.47 | 4.67 | 25.37 | 2.74 | 1.72 | 0.42 | 83.48 | 0.67 | 6.48 | 2.37E-05 | 2.01E-55 |
| Pass-by | 14.70 | 3.66 | 12.93 | 2.85 | 71.82 | 7.74 | 4.45 | 0.32 | 10.36 | 4.50E-17 | 2.97E+00 |

***b***

| | Participants | | SM Model | | State Model | | Movement Model | | $BF_{01}$ | | |
|---|---|---|---|---|---|---|---|---|---|---|---|
| *Trial type* | *M* | *SE* | *M* | *SE* | *M* | *SE* | *M* | *SE* | *Participants vs SM Model* | *Participants vs S Model* | *Participants vs M Model* |
| E-Indirect | 83.71 | 3.59 | 78.62 | 2.28 | 66.67 | 0.00 | 17.21 | 0.99 | 3.52 | 3.90E-05 | 1.62E-82 |
| E-Direct | 49.65 | 6.77 | 47.05 | 1.81 | 27.33 | 7.70 | 7.43 | 0.55 | 9.81 | 1.73E-07 | 2.51E-86 |
| W-Indirect | 9.23 | 3.08 | 7.76 | 1.38 | 29.63 | 3.70 | 77.24 | 1.19 | 0.86 | 5.56E+00 | 9.42E+00 |
| W-Direct | 29.73 | 6.08 | 41.14 | 4.34 | 0.52 | 0.30 | 89.12 | 0.58 | 7.26 | 2.14E-12 | 3.85E-45 |
| P-Indirect | 7.06 | 2.01 | 13.62 | 1.91 | 3.70 | 3.70 | 5.55 | 0.55 | 0.00 | 1.07E-32 | 3.51E-136 |
| P-Direct | 20.63 | 5.61 | 11.81 | 3.70 | 72.16 | 7.65 | 3.45 | 0.35 | 0.01 | 2.08E-102 | 4.12E-11 |

***c***

| | Participants | | SM Model | | State Model | | Movement Model | | $BF_{01}$ | | |
|---|---|---|---|---|---|---|---|---|---|---|---|
| *Message profile* | *M* | *SE* | *M* | *SE* | *M* | *SE* | *M* | *SE* | *Participants vs SM Model* | *Participants vs S Model* | *Participants vs M Model* |
| E-Forward | 18.59 | 1.24 | 17.46 | 0.80 | 6.14 | 1.73 | 0.43 | 0.03 | 8.40 | 1.12E-12 | 2.63E-27 |
| E-Backward | 15.30 | 1.21 | 14.58 | 0.50 | 4.01 | 1.13 | 0.42 | 0.02 | 9.06 | 3.88E-18 | 1.00E-31 |
| E-Left/right | 31.44 | 2.50 | 30.09 | 1.04 | 15.27 | 4.31 | 1.27 | 0.07 | 10.04 | 4.42E-04 | 5.91E-15 |
| W-Forward | 5.41 | 1.16 | 6.24 | 0.55 | 0.67 | 0.18 | 18.46 | 0.11 | 7.20 | 1.82E-05 | 2.27E-43 |
| W-Backward | 10.01 | 2.17 | 12.09 | 1.32 | 1.02 | 0.38 | 32.60 | 0.18 | 5.62 | 5.43E-05 | 4.08E-33 |
| W-Left/right | 6.91 | 1.42 | 8.21 | 0.85 | 0.99 | 0.25 | 46.13 | 0.15 | 5.96 | 4.52E-05 | 3.70E-235 |
| P-Forward | 4.26 | 1.15 | 3.63 | 0.67 | 17.39 | 2.00 | 0.16 | 0.02 | 10.06 | 1.07E-12 | 5.84E-01 |
| P-Backward | - | - | - | - | - | - | - | - | - | - | - |
| P-Left/right | 6.54 | 1.66 | 7.71 | 1.47 | 39.96 | 4.00 | 0.52 | 0.04 | 10.09 | 8.02E-23 | 1.91E+00 |

**Supplementary Table S1 | Summary of Behavioral Data and Computational Models with Bayesian Factor Comparison Across Different Message Types, Trial Types and Message profile for Dataset 1**. (a) Presents the results for the behavioral data, Surprise Model (SM), State Model, and Movement Model for three message types (Enter-exit, Wiggly, Pass-by) with mean percentages (M), standard errors (SE), and Bayesian factor comparison (BF01) between models for each message type. (b) Displays trial type results for indirect and direct conditions, including mean percentage, standard errors, and BF01values. (c) Details the message profile analysis for each experimental condition (E-Backward, E-Forward, E-Left/right, W-Backward, W-Forward, W-Left/right, P-Backward, P-Forward, P-Left/right), with participants' mean percentage, standard errors, and BF01 values for comparisons between the SM Model, State Model, and Movement Model.

***Data set 2***

***a***

| | Participants | | Surprise Model | | State Model | | Movement Model | | $BF_{01}$ | | |
|---|---|---|---|---|---|---|---|---|---|---|---|
| *Message type* | *M* | *SE* | *M* | *SE* | *M* | *SE* | *M* | *SE* | *Participants vs Surprise m* | *Participants vs S Model* | *Participants vs M Model* |
| Enter-exit | 54.95 | 6.6 | 55.24 | 1.95 | 25.48 | 7.28 | 9.65 | 0.42 | 11.12 | 1.96E-03 | 1.50E-08 |
| Wiggly | 26.65 | 5.91 | 27.55 | 3.86 | 0.07 | 0.07 | 86.02 | 0.39 | 10.95 | 8.14E-06 | 2.63E-30 |
| Pass-by | 18.41 | 4.95 | 17.2 | 3.11 | 74.45 | 7.27 | 4.33 | 0.33 | 10.95 | 2.48E-15 | 1.15E+00 |

***b***

| | Participants | | Surprise Model | | State Model | | Movement Model | | $BF_{01}$ | | |
|---|---|---|---|---|---|---|---|---|---|---|---|
| *Trial type* | *M* | *SE* | *M* | *SE* | *M* | *SE* | *M* | *SE* | *Participants vs Surprise m* | *Participants vs S Model* | *Participants vs M Model* |
| E-Indirect | 77.68 | 5.4 | 73.27 | 2.21 | 63.33 | 7.78 | 18.32 | 1.14 | 7.43 | 0.15 | 1.83E-31 |
| E-Direct | 48.93 | 7.18 | 49.75 | 2.04 | 26.16 | 7.47 | 7.01 | 0.44 | 11.13 | 1.51 | 2.29E-29 |
| W-Indirect | 13.14 | 4.8 | 13.02 | 2.77 | 3.33 | 3.33 | 70.51 | 0.97 | 6.51 | 2.63E-06 | 1.00E+01 |
| W-Direct | 30.73 | 6.46 | 31.98 | 4.44 | 0.04 | 0.04 | 90.74 | 0.46 | 10.94 | 1.42E-05 | 1.20E-19 |
| P-Indirect | 9.18 | 3.35 | 13.71 | 1.69 | 33.33 | 4.97 | 11.18 | 1.12 | 10.32 | 2.65E-19 | 1.01E-74 |
| P-Direct | 20.34 | 5.33 | 18.27 | 3.7 | 73.81 | 7.46 | 2.24 | 0.26 | 9.06 | 3.10E-59 | 1.68E-06 |

***c***

| | Participants | | Surprise Model | | State Model | | Movement Model | | $BF_{01}$ | | |
|---|---|---|---|---|---|---|---|---|---|---|---|
| *Message profile* | *M* | *SE* | *M* | *SE* | *M* | *SE* | *M* | *SE* | *Participants vs Surprise m* | *Participants vs S Model* | *Participants vs M Model* |
| E-Forward | 14.05 | 1.53 | 17.99 | 1.52 | 5.68 | 1.62 | 0.37 | 0.03 | 1.31 | 7.45E-04 | 7.77E-11 |
| E-Backward | 13.08 | 1.54 | 13.63 | 0.6 | 3.98 | 1.14 | 0.34 | 0.02 | 10.31 | 1.19E-08 | 2.96E-17 |
| E-Left/right | 28.65 | 3.64 | 26.91 | 0.85 | 15.18 | 4.34 | 0.97 | 0.06 | 10.15 | 0.04 | 7.79E-10 |
| W-Forward | 7.38 | 1.65 | 6.63 | 0.91 | 0.04 | 0.04 | 18.65 | 0.12 | 9.51 | 3.12E-06 | 4.29E-15 |
| W-Backward | 12.32 | 2.6 | 12.71 | 1.72 | 0.09 | 0.09 | 32.53 | 0.15 | 10.96 | 2.30E-06 | 6.55E-18 |
| W-Left/right | 8.15 | 1.75 | 9.07 | 1.27 | 0.06 | 0.06 | 46.6 | 0.18 | 9.3 | 1.02E-05 | 3.59E-136 |
| P-Forward | 4.5 | 1.23 | 3.89 | 0.73 | 18.69 | 1.89 | 0.07 | 0.01 | 10.42 | 3.30E-15 | 0.34 |
| P-Backward | - | - | - | - | - | - | - | - | - | - | - |
| P-Left/right | 7.36 | 2.28 | 9.16 | 1.52 | 44.85 | 4.11 | 0.46 | 0.04 | 9.75 | 1.08E-24 | 1.58 |

**Supplementary Table S2 | Summary of Behavioral Data and Computational Models with Bayesian Factor Comparison Across Different Message Types, Trial Types and Message profile for Dataset 2**. (a) Presents the results for the behavioral data, Surprise Model (SM), State Model, and Movement Model for three message types (Enter-exit, Wiggly, Pass-by) with mean percentages (M), standard errors (SE), and Bayesian factor comparison (BF01) between models for each message type. (b) Displays trial type results for indirect and direct conditions, including mean percentage, standard errors, and BF01values. (c) Details the message profile analysis for each experimental condition (E-Backward, E-Forward, E-Left/right, W-Backward, W-Forward, W-Left/right, P-Backward, P-Forward, P-Left/right), with participants' mean percentage, standard errors, and BF01 values for comparisons between the SM Model, State Model, and Movement Model.

***Data set 2 Predictive Accuracy of the Surprise Model across Different Samples***

***a***

| | Participants | | Surprise model, Sample1 pars | | $BF_{01}$ |
|---|---|---|---|---|---|
| *Message type* | *M* | *SE* | *M* | *SE* | *Participants vs Surprise m* |
| Enter-exit | 54.95 | 6.6 | 57.34 | 0.92 | 11.60 |
| Wiggly | 26.65 | 5.91 | 38.49 | 0.49 | 1.23 |
| Pass-by | 18.41 | 4.95 | 4.17 | 0.27 | 0.8 |

***b***

| | Participants | | Surprise model, Sample1 pars | | $BF_{01}$ |
|---|---|---|---|---|---|
| *Message profile* | *M* | *SE* | *M* | *SE* | *Participants vs Surprise m* |
| E-Forward | 14.05 | 1.53 | 11.43 | 0.24 | 3.86 |
| E-Backward | 13.08 | 1.54 | 12.38 | 0.22 | 11.1 |
| E-Left/right | 28.65 | 3.64 | 31.96 | 0.61 | 6.31 |
| W-Forward | 7.38 | 1.65 | 8.32 | 0.24 | 10.18 |
| W-Backward | 12.32 | 2.6 | 17.14 | 0.38 | 0.42 |
| W-Left/right | 8.15 | 1.75 | 14.33 | 0.4 | 0.004 |
| P-Forward | 4.5 | 1.23 | 0.84 | 0.09 | 0.93 |
| P-Backward | - | - | - | - | - |
| P-Left/right | 7.36 | 2.28 | 3.6 | 0.23 | 4.09 |

**Supplementary Table S3 | Predictive Accuracy of Surprise Model across Different Samples**

(a) Presents the results for the behavioral data and Surprise Model data (simulated for the second sample with the group estimated parameters of the first sample) for three message types (Enter-exit, Wiggly, Pass-by) with mean percentages (M), standard errors (SE), and Bayesian factor comparison (BF01) between models for each message type. (b) Displays Message profile results, including mean percentage, standard errors, and BF01values.
